# Similarity-Based Multimodal Regression

**DOI:** 10.1101/2022.04.13.488201

**Authors:** Andrew A. Chen, Sarah M. Weinstein, Azeez Adebimpe, Ruben C. Gur, Raquel E. Gur, Kathleen Ries Merikangas, Theodore D. Satterthwaite, Russell T. Shinohara, Haochang Shou

## Abstract

To better understand complex human phenotypes, large-scale studies have increasingly collected multiple data modalities across domains such as imaging, mobile health, and physical activity. The properties of each data type often differ substantially and require either separate analyses or extensive processing to obtain comparable features for a combined analysis. Multimodal data fusion enables certain analyses on matrix-valued and vector-valued data, but it generally cannot integrate modalities of different dimensions and data structures. For a single data modality, multivariate distance matrix regression provides a distance-based framework for regression accommodating a wide range of data types. However, no distancebased method exists to handle multiple complementary types of data. We propose a novel distance-based regression model, which we refer to as Similarity-based Multimodal Regression (SiMMR), that enables simultaneous regression of multiple modalities through their distance profiles. We demonstrate through simulation, imaging studies, and longitudinal mobile health analyses that our proposed method can detect associations in multimodal data of differing properties and dimensionalities, even with modest sample sizes. We perform experiments to evaluate several different test statistics and provide recommendations for applying our method across a broad range of scenarios.

## 1 Introduction

Complex health outcomes are understood as a byproduct of intricate biological pathways that are rarely captured in a single measurement. Advances in technology have enabled researchers to collect a large number of measurements on a single individual, spanning domains such as genomics, imaging, and physical activity. These individual data types are often called modalities, and the aggregation of several modalities on the same subject is called multimodal data. The availability of large multimodal data has increased considerably in the past decade, with studies such as the UK Biobank releasing multimodal data on roughly half a million individuals (Sudlow et al., 2015).

We focus on two large-scale multimodal studies, one collecting multimodal neuroimaging data and the other collecting a multitude of mobile health data. The Philadelphia Neurodevelopmental Cohort (PNC; Satterthwaite et al., 2014) consists of over 1,600 subjects with multimodal imaging including structural magnetic resonance imaging (MRI), functional MRI (fMRI), and diffusion tensor imaging (DTI). With the goal of understanding neurodevelopmental trajectories, studies have leveraged the PNC data to understand the effect of brain development on matrix-valued brain connectivity (Baum et al., 2020) and high-dimensional measures of cortical structures (Vandekar et al., 2015), among many other measures. For mobile health data, the National Institute of Mental Health (NIMH) Family Study of Affective Spectrum Disorders collects real-time data on over two hundred participants on their physical activity and emotional state through actigraphy and ecological momentary assessment (EMA) administered through mobile devices (Merikangas et al., 2014, 2019). Through the NIMH Family Study, researchers have identified differences among participants with mood disorders such as bipolar disorder in their patterns of sleep, mood, and physical activity (Merikangas et al., 2019). In both studies, there is a need for flexible methods to handle multimodal data, allowing further analyses while minimizing loss of information from the original data.

The emergence of these multimodal studies has driven methods for integration of multiple data modalities, often called multimodal data fusion. These methods vary considerably in their models and applications, but generally involve extensions of traditional multivariate analysis techniques such as independent component analysis, canonical correlation analysis, and singular value decomposition (Lahat et al., 2015). While these techniques work very well for certain types of data for which model constraints are satisfied, they are difficult to generalize to others. In neuroimaging for example, methods for integrating multiple modalities of functional imaging data may not directly apply to simultaneous analysis of structural and functional imaging. For more generally applicable models, several deep learning frameworks have been extended and enable prediction using multimodal data (Gao et al., 2020). However, these machine learning approaches are tailored for prediction tasks and are not suited for inference.

For analysis of data having arbitrary dimension and structure, distance-based and kernel-based methods provide inference through similarity metrics computed between subjects. Distance correlation and the Hilbert-Schmidt independence criterion are used for independence testing and are equivalent under certain choices of the distance and kernel (Sejdinovic et al., 2013; Shen and Vogelstein, 2020). Maximum mean discrepancy is used for two-sample kernel testing (Gretton et al., 2007) and permutational analysis of variance for multiple group distance-based tests (Anderson, 2001), with asymptotic properties recently investigated (Shinohara et al., 2020). For regression, multivariate distance matrix regression (MDMR; McArdle and Anderson, 2001) and kernel machine regression (KMM; Kwee et al., 2008) are both widely used and have been shown to be equivalent under certain conditions on the corresponding distance and kernel matrices (Pan, 2011). For multiple kernels computed on the same data, the microbiome regression-based kernel association test, an extension of KMM, allows for testing based on the minimum of *p*-values across kernels (Zhao et al., 2015). Another extension of KMM can incorporate multiple data modalities in regression on outcomes with distributions in the exponential family (Alam et al., 2021). To the best of our knowledge, none of these methods are designed to perform regression on multiple data modalities through their distances or kernels.

We propose a distance-based model for simultaneous regression of multimodal data of arbitrary types, which we call similarity-based multimodal regression (SiMMR). We develop two test statistics that are appropriate for different settings and compare them through simulation and applications to the PNC and NIMH Family Study data. We demonstrate that our test statistics outperform existing distance-based methods and provide high power for detection of associations across all data types considered. Our method introduces a novel framework for multimodal data fusion, which we demonstrate to be a flexible and powerful model for regression of multimodal data.

## 2 Methods

### 2.1 Multivariate distance matrix regression

We briefly review a standard distance-based method for regression of a single data modality called multivariate distance matrix regression (MDMR; McArdle and Anderson, 2001; Anderson, 2001; Schork and Zapala, 2012). Let (Ω, *d*) be a semimetric space and *Y* be a random object taking values in Ω. Suppose we observe independent draws *y*_*i*_ for each subject *i* = 1, 2, …, *n* and let *D* = (*d*_*ij*_)_*n×n*_ denote the sample dissimilarity matrices where *d*_*ij*_ = *d*(*y*_*i*_, *y*_*j*_). Define the doubly-centered dissimilarity matrix *G* = (*I* − **11**^*T*^)*A*_*k*_(*I* − **11**^*T*^) where 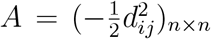 Let *X* be an *n* × *p* design matrix with corresponding projection matrix *H* = *X*(*X*^*T*^ *X*)^−1^*X*^*T*^.

MDMR tests for an association of *Y* and *X* via the psuedo-F statistic

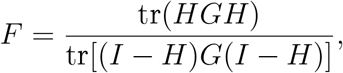

where statistical significance is typically evaluated through permutation. With a univariate outcome and Euclidean distance, it is equivalent to the standard regression *F* -statistic if appropriate degrees of freedom are accomodated (McArdle and Anderson, 2001). For testing subsets of covariates, the numerator can be replaced by tr[(*H* − *H*_*r*_)*G*(*H* − *H*_*r*_)] where *H*_*r*_ is the projection matrix from the reduced model (Reiss et al., 2010).

Recent papers investigate the asymptotic null distribution of the MDMR test statistic, deriving distributions based on *χ*^2^ random variables weighted by the eigenvalues of *G*. McAr-tor et al. (2017) reformulate the statistic in terms of multidimensional scaling (MDS) scores and derive the asymptotic distribution as a quotient of weighted sums of central *χ*^2^ random variables by making assumptions about the distribution of MDS scores. By assuming matrix normal error and limiting their scope to Euclidean and Mahalanobis distances, Li et al. (2019) find the distribution to be a weighted quotient of non-central *χ*^2^ random variables. For distance-based analysis of variance, Shinohara et al. (2020) represent the psuedo-F statistic as a *U* -statistic to identify the limiting distribution as a weighted sum of central *χ*^2^ random variables. Shi et al. (2021) adapt results from kernel-based testing to derive the asymptotic null distribution as a weighted sum of non-central *χ*^2^ random variables, making no assumptions about the error structure or limitations on the distance function.

### 2.2 Similiarity-based multimodal regression model

Let (Ω_1_, *d*_1_), (Ω_2_, *d*_2_), …, (Ω_*m*_, *d*_*m*_) be semimetric spaces. We consider random objects *Y*_1_, *Y*_2_, …, *Y*_*m*_ taking values in Ω_1_, Ω_2_, …, Ω_*m*_ respectively and a vector-valued random variable for covariates *X*. Suppose we observe independent draws {*y*_1*i*_, *y*_2*i*_, …, *y*_*mi*_} as the multimodal outcome and *x*_*i*_ as the corresponding vector of covariates of dimension length *p* for each subject *i* = 1, 2, …, *n*. Let *D*_*k*_ = (*d*_*kij*_)_*n×n*_ denote sample dissimilarity matrices defined based on appropriately chosen distance metrics for each individual data modality where *d*_*kij*_ = *d*_*k*_(*y*_*ki*_, *y*_*kj*_) for *k* = 1, 2, …, *m*.

Our goal is to assess the joint assocation between *Y*_1_, *Y*_2_, …, *Y*_*m*_ and *X* through their respective dissimilarity matrices *D*_1_, *D*_2_, …, *D*_*m*_. Define weighted doubly-centered dissimilarity matrices *G*_1_, *G*_2_, …, *G*_*m*_ where *G*_*k*_ = *w*_*k*_(*I* − **11**^*T*^)*A*_*k*_(*I* − **11**^*T*^) and 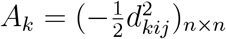 In our analyses, the weights *w*_*k*_ > 0 are chosen as the largest eigenvalue of *G*_*k*_ following recommendations from previous literature on integration of multiple distance matrices (Abdi et al., 2005). Weights based on other properties of *G*_*k*_ may alternatively be selected, or weights can be chosen to place particular emphasis on certain modalities.

Our model tests for an association between these weighted doubly-centered dissimilarity matrices and the covariates of interest. These *G*_*k*_ admit the decompositions 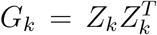 where *Z*_*k*_ = [*z*_*k1*_ *z*_*k2*_ … *z*_*kn*_]^*T*^ are the *n*-dimensional classical multidimensional scaling (cMDS) scores (McArdle and Anderson, 2001). Note that for non-Euclidean distances, *G*_*k*_ is not guaranteed to be positive semidefinite and cMDS scores may include imaginary values. One solution is to discard imaginary cMDS axes; however, McArdle and Anderson (2001) show that this might lead to conservative tests. The recommended solution is to add a constant to off-diagonal elements of each distance matrix prior to computation of cMDS, which has a solution derived in Cailliez (1983) and recently applied in the formulation of partial distance correlation (Székely and Rizzo, 2014).

Let *Z* = [*Z*_1_ *Z*_2_ … *Z*_*m*_] denote the *n* × *mn* matrix of concatenated cMDS scores; this concatenation was first proposed in Faraway (2014). We propose similarity-based multimodal regression (SiMMR) as the multivariate regression model

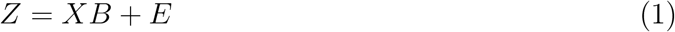

where *X* = [*x*_1_ *x*_2_ … *x*_*n*_]^*T*^ is the *n* × *p* design matrix, *B* is a *p* × *mn* matrix of regression coefficients and *E* is an *n*×*mn* error matrix. Our null hypothesis is *H*_0_ : *B* = **0**, corresponding to no association between *Y* and *X*,

### 2.3 SiMMR-D and SiMMR-PC test statistics

We propose two statistics to test the null hypothesis that a subset of covariates has no association with the multimodal outcome *Y*. Let *B* = (*B*_1_ *B*_2_)^*T*^ *B* where *B*_2_ are the covariates of interest. Our goal is to test the null hypothesis *H*_0_ : *B*_2_ = **0** against *H*_*a*_ : *B*_2_ = **0**. Standard test statistics for multivariate regression require *Z* to be full-rank and compare the sum of squares and cross products (SSCP) matrices of the hypothesis 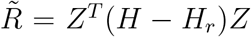 and the SSCP error matrix 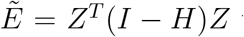 where *H* = *X*(*X*^*T*^ *X*)^−1^*X*^*T*^ and *H*_*r*_ is the hat matrix from the reduced model. In our case, *Z* is rank deficient since *mn > n* for *m >* 1. To perform regression in this high-dimensional setting, we propose two alternative approaches. First, we adapt the Dempster trace (Dempster, 1958) to our setting, which we denote by *T*_*D*_, the SiMMR Dempster trace (SiMMR-D). Directly applying the original formulation of Dempster trace, SiMMR-D is the ratio between the traces of the SSCP matrices,

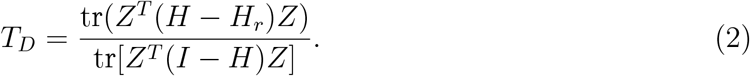

Using the idempotency of *H* and the cyclical property of the trace, we can rewrite this as

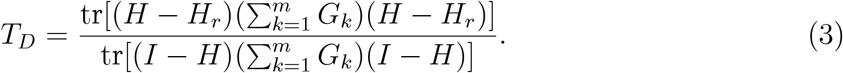

This equality shows that SiMMR-D is equivalent to performing MDMR using the sum of the dissimilarity matrices, which connects SiMMR-D directly to this classic distance-based regression framework. Theoretical results derived for MDMR thus apply directly to our method, which include the asymptotic null distribution of the MDMR psuedo-F statistic previously discussed in Section 2.1.

The SiMMR-D test statistic is a natural extension of MDMR, but it discards crossproduct terms in the SSCP matrices that capture the correlations among modalities. As an alternative solution, we propose another test statistic that leverages the correlations of the cMDS scores. We first address the rank deficiency of the SSCP matrices by performing dimension reduction on the cMDS scores using principal component analysis (PCA). We then construct Pillai’s trace from the first *K* PC scores represented in the *n* × *K* matrix *W*. This test statistic, which we denote by *T*_*PC*_ and we call SiMMR principal components (SiMMR-PC(*K*)), is defined through the corresponding *K* × *K* SSCP matrices 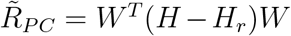 and 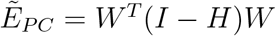 as

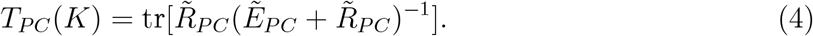

For the design matrix of dimensions *n* × *p, T*_*PC*_(*K*) is defined for *K* < *n* − *p* − 1. We investigate the choice of *K* through simulation and applications in Sections 4.1 and 4.3.

SiMMR-PC performs inference based on the cMDS scores of each modality, which are not unique since any orthogonal rotation of the optimal scores is also optimal (see for example, page 396 in Mardia et al. (1979)). Orthogonal rotation of each cMDS solution can be represented as a rotation of the concatenated cMDS scores. However, owing to rotational invariance of the Pillai’s trace test statistic (Langsrud, 2004), any set of cMDS solutions yields the same *T*_*PC*_ test statistic.

Testing for both SiMMR-D and SiMMR-PC statistics proceeds via permutation. The permutational null distribution is generated by permuting the rows of the design matrix and computing the chosen SiMMR test statistic. The *p*-value for the permutation test is then the proportion of permuted test statistics less than the test statistic computed using the original design matrix. Throughout our analyses, we perform 999 permutations for computation of each SiMMR *p*-value.

## 3 Simulation study

We evaluate the efficacy of our proposed SiMMR methodology through a simulation study with varying sample size, number of features, number of modalities, and correlation structure. We compare our method to MDMR and multivariate multiple regression (MMR) where applicable.

### 3.1 Data generation

We simulate a multimodal dataset with correlations within and between modalities through the following setup. Let *N* be the simulation sample size, *M* be the number of modalities, and *Q* be the number of features per modality. Let *Y* = (*Y*_1_, *Y*_2_, …, *Y*_*M*_) denote the full *MQ*-dimensional vector of multimodal data and *x* ∼ Binomial(0.5) denote a simulated binary covariate.

We draw the *N* multimodal observations from a multivariate normal distribution with correlation structure Σ of dimension *MQ* × *MQ* where the covariate shifts the observations in directions of the eigenvectors of Σ. For a covariate effect of rank *L*, let 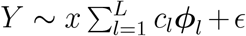 where ∈ ∼ *N* (**0**, Σ) where Σ is the chosen correlation structure and ***ϕ***_*l*_, *l* = 1, 2, …, *MQ* are the eigenvectors of Σ.

We propose three simulation settings by modifying the correlation structure. In our first scenario, Σ is an exchangeable correlation matrix with parameter *ρ* = 0.25 to simulate multimodal data that have low correlations within modality and between modalities. Our second scenario uses an exchangeable correlation structure with *ρ* = 0.75 so that the simulated multimodal data has high correlations. The third scenario has Σ instead as a first-order autoregressive structure, or AR(1), with parameter *τ* = 0.9. The AR(1) structure yields correlations that are generally higher within modality than between modalties.

In each scenario, we vary the rank and magnitude of the covariate effect. We choose *L* as 1, ⌊*M Q/*4⌋, ⌊*M Q/*2⌋, and *MQ* to provide simulation settings with varying complexity of covariate effects. The contribution of each eigenvector of Σ to the covariate effect varies across settings to ensure that the strength of the effect remains similar. In particular, *L* = 1 has *c*_1_ = 3, *L* = *M Q/*4 has *c*_1_ = *c*_2_ = … = *c*_*MQ/*4_ = 0.7, *L* = *M Q/*4 has *c*_1_ = *c*_2_ = … = *c*_*MQ/*2_ = 0.7, and *L* = *MQ* has *c*_1_ = *c*_2_ = … = *c*_*MQ*_ = 0.15.

We compare SiMMR to two competing methods: multivariate multiple regression (MMR) and multivariate distance matrix regression applied to each modality correcting for multiple comparisons using Bonferroni correction (MC-MDMR). For simulation settings where *MQ* < *N* −2, we perform MMR by computing Pillai’s trace based on the simulated data *Y*. SiMMR-PC is computed with the number of PCs *K* ranging from 2 to 25. For SiMMR-PC with *K* ≥ *N* − 2, all PCs are included.

### 3.2 Simulation results

#### Type I error is well-controlled across simulation settings and SiMMR test statistics

In Table 1, we display the type I error and power of MMR, MC-MDMR, SiMMR-D, and SiMMR-PC(3) across simulation settings. **Supplementary Table 1** shows results for AR(1) correlation settings while also including SiMMR-PC(10) and SiMMR-PC(15). We find that the type I error rates for SiMMR test statistics are well-controlled across simulation settings, but MC-MDMR is overly conservative at high numbers of modalities *M*, especially in the exchangable correlation settings. Across correlation structures at sample size 100 and 10 modalities, MMR has a slightly conservative type I error.

**Table 1:**
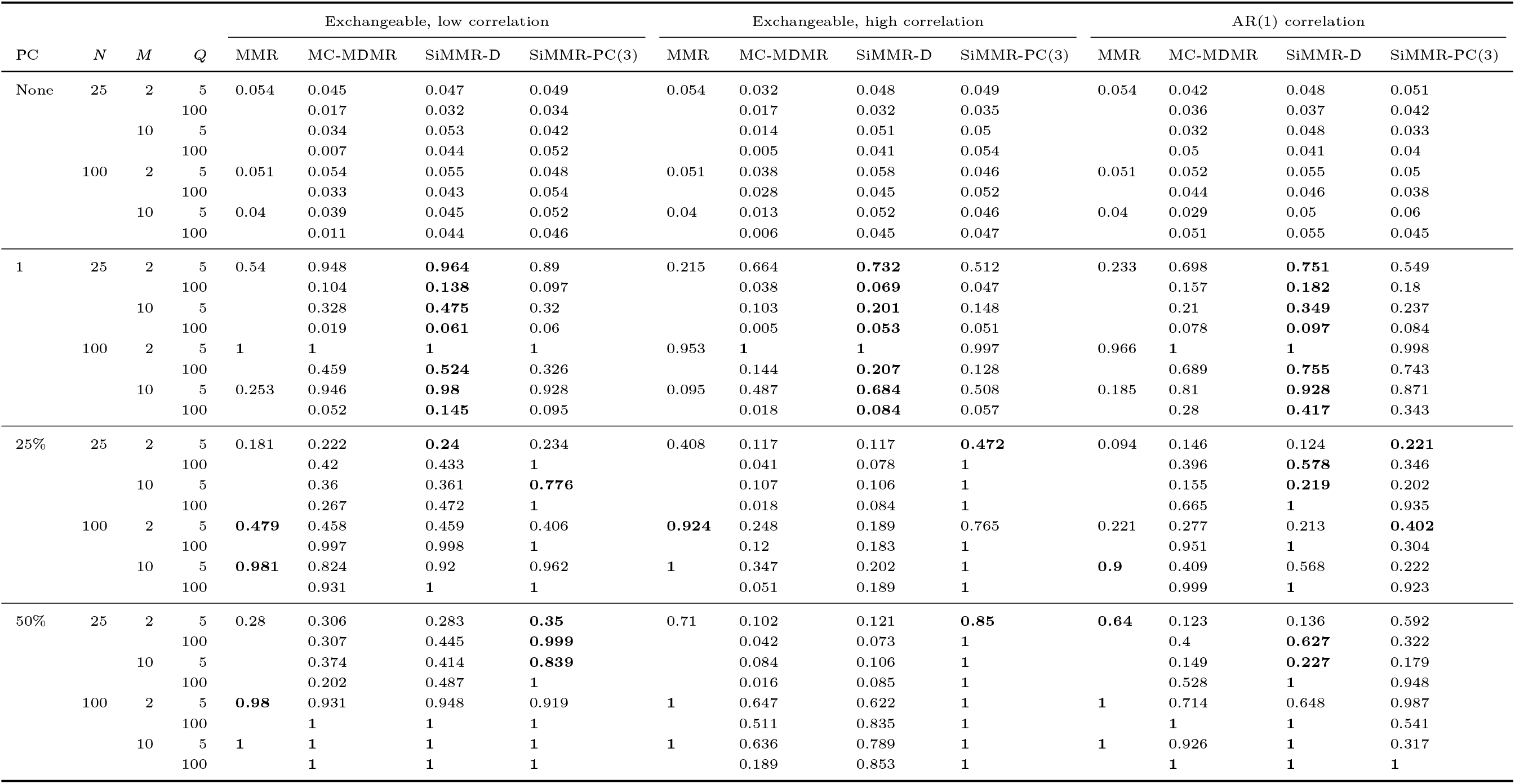
Simulation results across varying sample size, number of features, covariate effect, and correlation structure. Rejection rates are shown across varying number of subjects (N), number of modalities (M), number of features per modality (Q), and number of principal components included in the binary covariate effect (PC). The highest power among tests within each simulation setting is bolded.

#### SiMMR outperforms MC-MDMR across simulation settings

For covariate effects in the first PC direction, SiMMR-D shows equal or greater power than MC-MDMR and MMR across all simulations. For more complex covariate effects, we observe that SiMMR-PC(3) yields greater power than MC-MDMR across the exchangeable correlation settings with especially large differences in the high correlation setting. **Supplementary Table 1** shows that either SiMMR-PC(10) or SiMMR-PC(15) outperform MC-MDMR across all AR(1) correlation settings. Fig. 2 and **Supplementary Fig. 1** compare test statistics for settings with two modalities and show that these results hold across number of features *Q* per modality.

**Figure 1:**
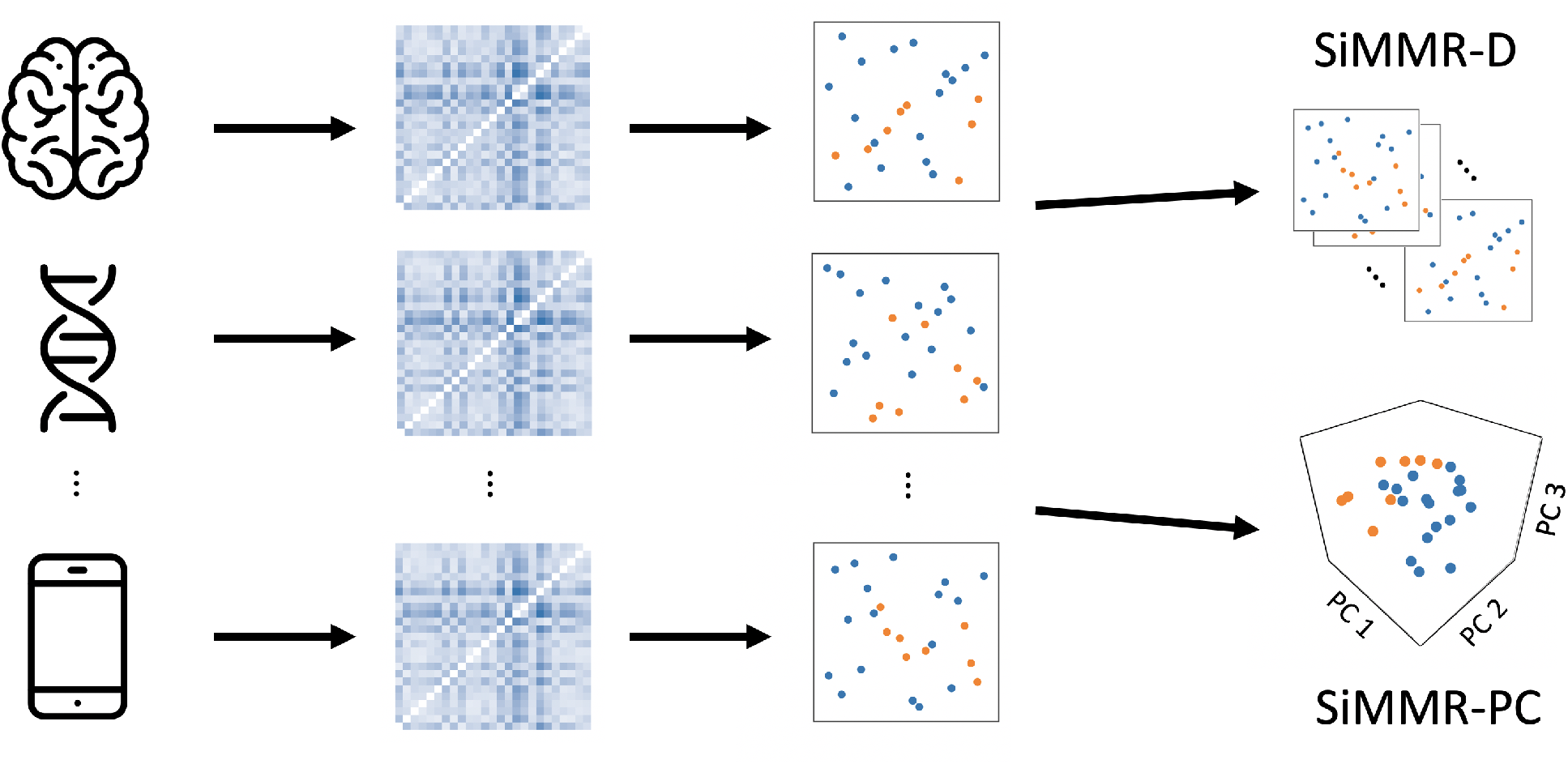
Illustration of similarity-based multimodal regression. In SiMMR, distance matrices are computed separately on each modality, followed by representation in Euclidean space via classical multidimensional scaling (cMDS). SiMMR then concatenates these cMDS coordinates and performs inference using either Dempster’s trace (SiMMR-D) or Pillai’s trace after dimension reduction using principal components (SiMMR-PC).

**Figure 2:**
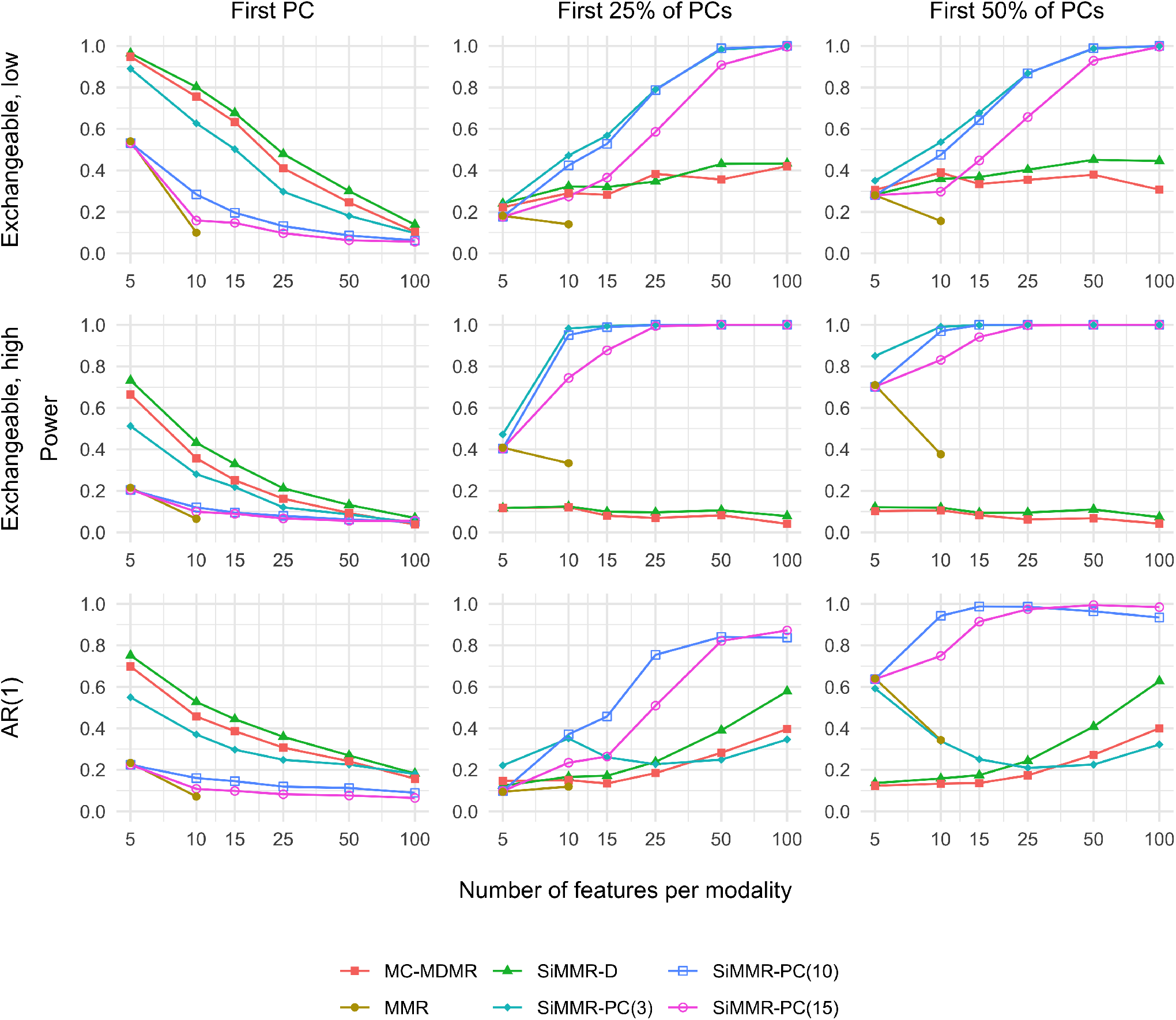
Power results in simulations with exchangeable and AR(1) correlation structures for a sample size of 25. Each trace represents a different test statistic. Different simulation settings are distinguished by correlation structure across rows and by rank of the binary covariate effect across columns. Exchangeable refers to an exchangeable correlation structure with low or high correlation and AR(1) refers to a first-order autoregressive structure. Abbreviations: MDMR, multivariate distance matrix regression; MC-MDMR, multiple MDMR statistics after Bonferroni correction; MMR, multivariate multiple regression using Pillai’s trace.

#### SiMMR yields comparable or higher power than MMR

In settings with *MQ < N* − 2, we can apply MMR and compare the power results to SiMMR. At a sample size of 25, we find that SiMMR-PC yields similar or greater power than MMR across all settings. SiMMR-PC(10) and SiMMR-PC(15) show particularly higher power in exchangeable, high correlation and AR(1) correlation settings (**Supplementary Table 1**). Across settings with a sample size of 100, **Supplementary Table 1** and **Supplementary Fig. 1** show that SiMMR-PC(3) has comparable or lesser power than MMR but SiMMR-PC(10) and SiMMR-PC(15) yield equal or greater power than MMR.

#### Relative performance of SiMMR-D and SiMMR-PC depends on correlation structure and covariate effect

In settings with a covariate effect in the first PC direction, Table 1 shows that SiMMR-D yields equal or higher power than SiMMR-PC and **Supplementary Figures 2 and 3** show that this relationship holds across SiMMR-PC test statistics for 2 through 25 PCs. For exchangeable correlation structures and covariate effects in 25% or 50% of the PC directions, SiMMR-PC outperforms across sufficiently low number of PCs in the low correlation setting and across all choices in the high correlation setting. For settings with an AR(1) correlation structure, the performance of SiMMR-PC relative to SiMMR-D depends on the complexity of the covariate effect and number of features per modality. SiMMR-PC with large numbers of PCs performs better in settings with more complex covariate effects and larger number of features. **Supplementary Table 1** numerically compares SiMMR-PC test statistics for AR(1) correlation settings and shows that SiMMR-PC(3) performs the best for covariate effects with 25% of PCs and 5 features per modality, but SiMMR-PC(10) and SiMMR-PC(15) outperform in other settings.

## 4 Data applications

We apply SiMMR to two studies with novel and distinct types of multimodal data. Our first application involves neuroimaging data from the Philadelphia Neurodevelopmental Cohort (PNC; Satterthwaite et al., 2014), where we are interested in testing for age-related changes in brain connectivity and cortical structure. Our second application uses mobile health data from the National Institute of Mental Health (NIMH) Family Study of Affective Spectrum Disorders (Merikangas et al., 2014), where we test for differences in mood and physical activity measures among subjects with mood disorders. Our SiMMR applications involve matrix-valued structural and functional connectivity from the PNC, vector-valued cortical thickness and sulcal depth measurements from the PNC, and time series observations of mobile health data from the NIMH Family Study of Affective Spectrum Disorders.

### 4.1 Philadelphia Neurodevelopmental Cohort

We apply the SiMMR methodology to two multimodal neuroimaging datasets collected as part of the PNC (Satterthwaite et al., 2014). All participants, or their parent or guardian, provided informed consent, and minors provided assent. The study was approved by the institutional review boards of both the University of Pennsylvania and the Children’s Hospital of Philadelphia. The PNC includes 9,498 subjects between the ages of 8 and 23. Multimodal imaging was acquired on a subset of 1,601 subjects using a Siemens TIM Trio 3-T scanner with a 32-channel head coil and the same imaging sequences and parameters for every subject. Included participants in the PNC were medically healthy, were not taking psychoactive medication, and passed strict quality-assurance procedures for their imaging.

### 4.2 Image acquisition and preprocessing

In this study, we first examine a set of subjects from the PNC with structural connectivity, resting-state functional connectivity, and n-back functional connectivity measurements. Acquisition and preprocessing for this sample has previously been discussed in Baum et al. (2020). In summary, 727 participants are included after strict quality assurance procedures (demographic details in Table 2 under “PNC Connectivity”). Structural connectivity is calculated from diffusion-weighted imaging using probabilistic tractography. Entries of each subject’s structural connectivity (SC) matrices are computed as the number of probabilistic streamlines connecting each pair of 400 brain regions, normalized by the total edge weight across all network connections. Functional connectivity matrices are computed separately for functional magnetic resonance imaging (fMRI) acquired while the participant is at rest (rsFC) and during the n-back task (n-back FC). Functional connectivity between each pair of the 400 brain regions is computed as the Pearson correlation coefficient between the mean regional blood-oxygen-level-dependent (BOLD) time series.

**Table 2:** Demographics of the imaging and mobile health datasets. Age, sex, and diagnosis status (applicable only to the NIMH Family Study) are shown for all subjects included in this study. The two PNC datasets are different subsets of the full PNC dataset with some overlap. Abbreviations: PNC, Philadelphia Neurodevelopmental Cohort; FC, functional connectivity; SC, structural connectivity; CT, cortical thickness; SD, sulcal depth; NIMH, National Institute of Mental Health; MDD, major depressive disorder.

|  | PNC Connectivity | PNC Cortical Structure | NIMH Family Study |
| --- | --- | --- | --- |
| Number of subjects | 727 | 912 | 77 |
| Age, mean(SD) | 15.88 (3.23) | 14.80 (3.50) | 49.31 (17.73) |
| Male, n(%) | 307 (42.2) | 415 (45.5) | 31 (40.3) |
| Diagnosis, n(%) |  |  |  |
| Healthy Control |  |  | 34 (44.2) |
| MDD |  |  | 26 (33.8) |
| Bipolar Type I |  |  | 6 ( 7.8) |
| Bipolar Type II |  |  | 11 (14.3) |

Our second PNC application uses a subset of 912 PNC subjects with high-quality cortical thickness (CT) and sulcal depth (SD) measurements computed from T1-weighted images (demographic details in Table 2 under “PNC Cortical Structure”). The acquisition and preprocessing for these data was originally described in Vandekar et al. (2015) and in subsequent studies (Vandekar et al., 2016; Weinstein et al., 2021). Cortical reconstruction of the T1-weighted structural images was completed using FreeSurfer (version 5.3). These cortical measurements were resampled to the fsaverage5 atlas, which has 10242 vertices in each brain hemisphere. Cortical thickness was computed as the minimum distance between pial and white matter surfaces (Dale et al., 1999) and sulcal depth as the height of gyri (Fischl et al., 1999).

#### 4.2.1 Application of SiMMR

Using SC and FC matrices from the PNC, Baum et al. (2020) demonstrated neurodevelopmental changes in a coupling metric computed between SC and n-back FC while separately examining coupling between SC and rsFC. In our application, we incorporate all three modalities and apply SiMMR to determine if there is an association between SC, n-back FC, and rsFC jointly with age while controlling for sex and relevant quality metrics. The quality metrics are identical to those in Baum et al. (2020), which includes mean relative framewise displacements calculated on the resting-state and n-back fMRI scans and mean relative displacement from interspersed volumes with a b value of 0 s/mm^2^ calculated from the diffusion-weighted images. We choose to compute dissimilarities between connectivity matrices using the Frobenius distance.

Previous studies of the PNC have demonstrated neurodevelopmental changes in cortical thickness (Vandekar et al., 2015) and the coupling between cortical thickness and sulcal depth (Vandekar et al., 2016). We apply SiMMR to the cortical structure data for regression on age while controlling for sex. We construct dissimilarity matrices based on the Euclidean distance.

To assess power in these applications, we use a resampling approach to evaluate rejection of the null hypothesis in smaller sample sizes. We draw 1000 subsamples without replacement and compute SiMMR *p*-values for each subsample. The power is then calculated as the proportion of *p*-values with value less than our nominal type I error rate of 0.05.

#### 4.2.2 Results

In PNC structural and functional connectivity data, Fig. 3a shows that SiMMR-D has high power for detecting age-related changes in connectivity, achieving 82% power at a sample size of 40. Comparing this multimodal analysis to unimodal analyses, only MDMR on structural connectivity achieves similar power with 74.2% power at the same sample size and combining all three unimodal analyses via MC-MDMR yields 71.6% power. These results suggest that multimodal structural and functional analysis via SiMMR is better able to detect changes in brain connectivity than any unimodal analysis or the combination of unimodal analyses.

For changes of cortical structure during brain development, joint analysis of cortical thickness and sulcal depth does not outperform unimodal analysis of cortical thickness. MDMR on cortical thickness has 83.3% power to detect an age association at a sample size of 20, whereas MC-MDMR and SiMMR-D only have 75.9% and 72.9% power respectively. At the same sample size, MDMR on sulcal depth only has 13% power. These results are consistent with a previous study finding age-related changes in cortical thickness using the PNC study (Vandekar et al., 2015). We find that age-related changes in sulcal depth require a larger sample size to detect, which leads inclusion of sulcal depth to reduce power in multimodal analyses. Our results do not contradict previous reports of age-related patterns in the coupling between cortical thickness and sulcal depth (Vandekar et al., 2016); however, this relationship does not drive higher power for detection of age when jointly analyzing the two modalities.

### 4.3 NIMH Family Study

We also apply SiMMR to mobile health data collected as part of the National Institute of Mental Health (NIMH) Family Study of Affective Spectrum Disorders, an observational cohort study of subjects recruited from the greater Washington, DC, metropolitan area (Merikangas et al., 2014). All participants provided informed consent, and the study was approved by the Combined Neuroscience Institutional Review Board at the National Institutes of Health. Each participant was evaluated for mental disorders via a comprehensive semistructured diagnostic interview, with mental disorders defined by Diagnostic and Statistical Manual for Mental Disorders, IV^th^ Edition (DSM-IV) criteria. Further details on the recruitment and exclusion criteria can be found in Merikangas et al. (2014).

#### 4.3.1 Mobile health data preprocessing

For our study, we examine the subset of 384 participants with actigraphy and ecological momentary assessment (EMA) data. Physical activity data was collected via accelerometers (Actiwatch Spectrum, Philips Respironics, Murrysville, PA, USA), which produced activity counts for every minute of the day measured via movement-related voltage signals recorded by the accelerometer. Participants completed EMA four times a day approximately four hours apart through a smartphone during the same two-week assessment period when accelerometry data are being collected. For our analyses, we include self-reported mood variables in EMA, which consist of 7-point Likert scales to measure the degree to which participants felt active, anxious, energetic, sad, distracted, and irritable. Further details on the actigraphy, EMA data collection, and activity data processing were presented in Johns et al. (2019), Lamers et al. (2018), Merikangas et al. (2019), and Shou et al. (2017).

For our analysis, we include 77 subjects with at least one week of data. We summarize the activity and EMA time series by averaging time points within each weekday across two weeks. The time points consist of 1440 minutes per day for activity data and 4 times per day for EMA data. For time points with missing data for either week, we use the single available measurement. We exclude subjects from our analyses who are missing both measurements for any time point of EMA data. Our mobile health dataset consists of 77 participants with 10080 activity time points and 28 EMA time points for each of the six EMA variables. Demographic details are available in Table 2.

#### 4.3.2 Application of SiMMR

A previous study used mobile health data from the NIMH Family Study to understand the joint relationship between physical activity and mood, and to identify diagnosis-related differences in the association between activity, energy, mood, and sleep (Merikangas et al., 2019). This study summarized the activity data into four bins per day to align with the EMA measurements, which only included measures of sadness and energy. We use SiMMR to identify diagnosis effects while leveraging the full activity time series as well as all six EMA variables. For each modality, we calculate dissimilarity matrices using the Euclidean distance between time series. Our analysis is performed by using SiMMR to simultaneously regress activity and the EMA variables on diagnosis status while controlling for age and sex. To assess power in this application, we use the resampling approach previously described in Section 4.2.1.

#### 4.3.3 Results

Fig. 3c demonstrates that SiMMR-PC can detect diagnosis-related changes jointly among physical activity and EMA measurements of mood with high power at larger sample sizes. Multimodal analysis using SiMMR-PC(3) has 83.2% power at sample size 50, which far outperforms unimodal MDMR analyses where the highest power is 57.1% from MDMR on EMA-measured feelings of irritability. EMA-measured feelings of anxiety and sadness and measures of physical activity as assessed by accelerometers also show associations with diagnosis, having 56.6%, 38.6%, and 25.5% power respectively at sample size 50. Other unimodal analyses show notably less power across sample sizes considered. Combining these unimodal analyses via MC-MDMR yields low power as a result with MC-MDMR having 26.2% power for the same number of subjects. These observations suggest that joint analysis of physical activity and mood measurements using SiMMR can identify diagnosis-related changes more effectively than use of existing methodologies for unimodal or combined unimodal analyses.

**Figure 3:**
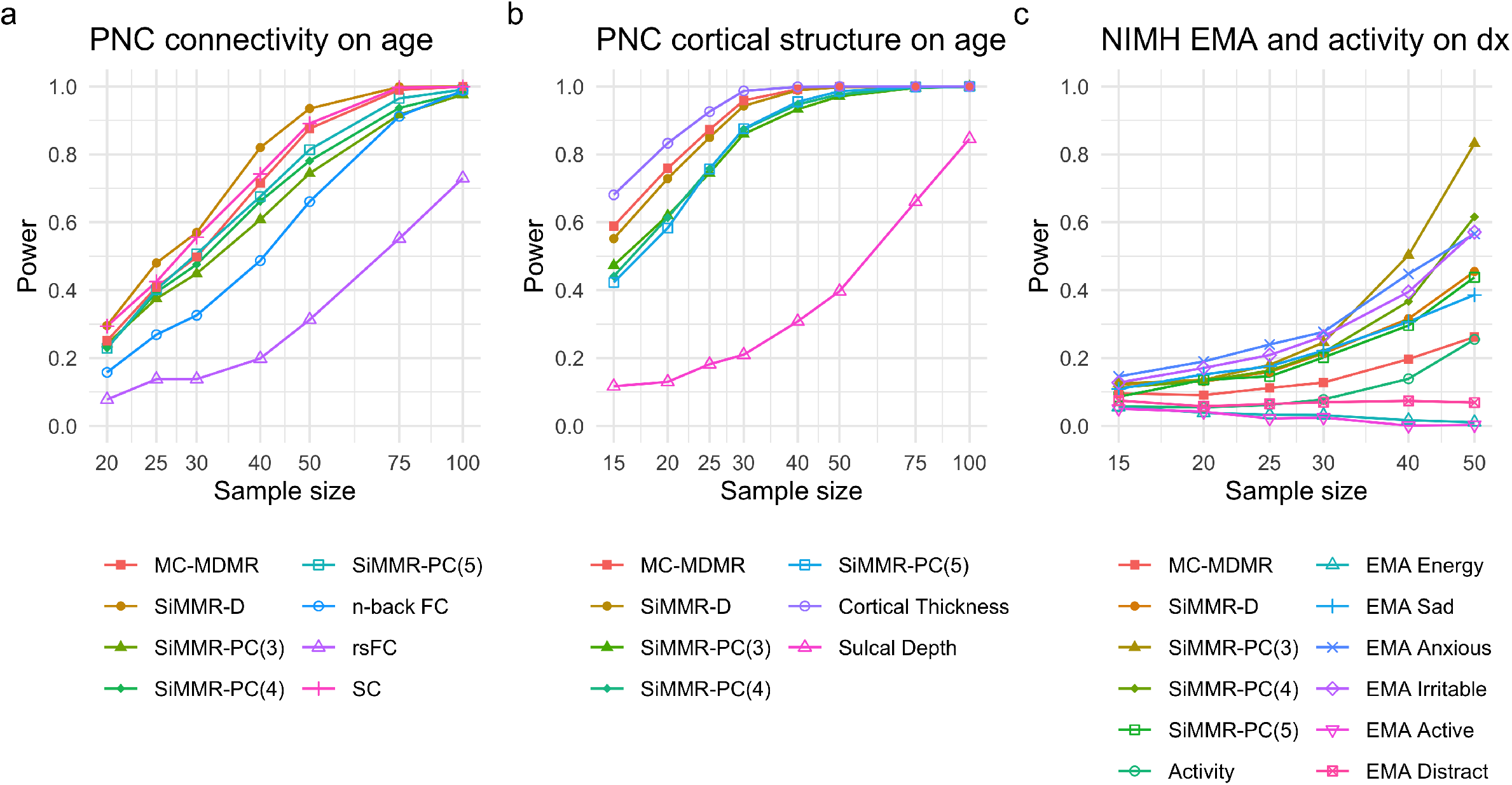
Power results in applications of SiMMR to imaging and mobile health data. Each trace represents a different test statistic. Power curves for individual modalities are obtained through multivariate distance matrix regression (MDMR). Abbreviations: PNC, Philadelphia Neurodevelopmental Cohort; FC, functional connectivity; rsFC, resting-state functional connectivity; SC, structural connectivity; EMA, ecological momentary assessment; MC-MDMR, multiple MDMR statistics after Bonferroni correction; MMR, multivariate multiple regression using Pillai’s trace.

### 4.4 Selection of SiMMR-PC

In our applications, selection of the number of PCs included in SiMMR-PC is important to detect associations of interest. Fig. 4a shows that in our PNC applications, SiMMR-D outperforms SiMMR-PC across all numbers of PCs considered. However, SiMMR-PC(3) and SiMMR-PC(4) have higher power for detection of diagnosis than SiMMR-D at higher sample sizes in the NIMH Family Study application. To investigate possible explanations behind such observations, we compute the distance correlation (Székely et al., 2007) between each modality using data from all subjects. Fig. 4b shows that the distance correlations in the PNC applications are relatively low (*<* 0.40) and the distance correlations among certain EMA measurements in the NIMH Family Study are considerably higher (> 0.60 for certain pairs of modalities). Fig. 4c further shows that the percent of variation explained by the first few PCs across resamplings of 50 subjects is considerably higher in the NIMH Family Study application. These findings demonstrate that distance correlation and scree plots from PCA can inform when to use SiMMR-PC and how to select the number of PCs included. Based on our observations, we suggest use of SiMMR-PC in applications with high distance correlation among modalities and choosing PCs that explain a large portion of the variation among MDS scores.

**Figure 4:**
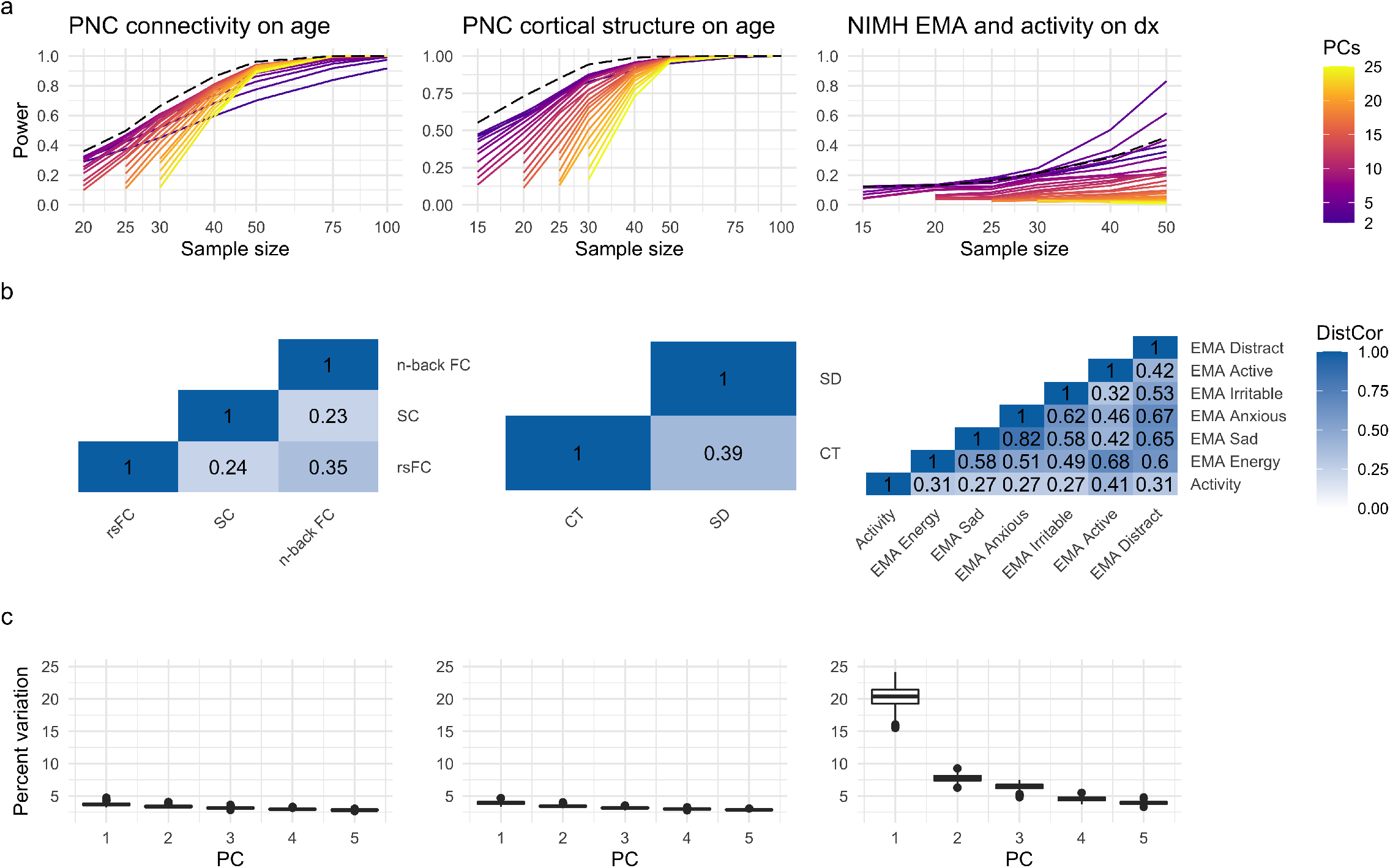
SiMMR-PC results across number of PCs and related exploratory analyses in real data applications. (a) shows the power of SiMMR-PC test statistics across number of PCs compared to SiMMR-D (dashed line). (b) displays the distance correlation (DistCor) among modalities in each application using the full sample. (c) shows the percent of variation explained by PCs across the 1000 resamplings of size 50 in each application. Abbreviations: PNC, Philadelphia Neurodevelopmental Cohort; EMA, ecological momentary assessment.

## 5 Discussion

The emergence of technology and organized efforts for collection of multiple types of health data provides a great opportunity to jointly examine associations between multimodal assessments and health outcomes. To integrate and perform inference in multimodal settings, we develop a flexible distance-based testing framework called SiMMR, which can incorporate data from arbitrary semimetric spaces. We demonstrate in simulation and real data that our test statistics can identify associations with relatively small sample sizes and across a wide range of data structures. We propose two alternative test statistics that provide higher power in certain settings, generally outperforming existing distance-based methods.

We find that relative performance of SiMMR versus unimodal analysis depend on the included modalities and their correlation. In our simulations and applications, the benefit of performing SiMMR is limited when modalities show lesser correlation. We also observe through our application to PNC cortical structure that modalities with weak associations can reduce power of a multimodal analysis, even when the correlation between modalities is known to be important (Vandekar et al., 2016). Our results emphasize that knowledge about the data structure should inform whether application of SiMMR is appropriate and the choice of modalities to include.

Comparing SiMMR test statistics, SiMMR-PC generally outperforms SiMMR-D when the correlation among modalities is high and the effect of interest is sufficiently complex. For selection among SiMMR-PC statistics, we found that the first three or four PCs provided optimal power across most of our settings, with additional PCs needed in simulations with complex correlation structures. We used standard scree plots to choose among PCs that explain the most variation; however, other investigations have suggested that PCs explaining less variation may be more closely associated with outcome measures (Liu et al., 2020). While our choices of SiMMR-PC statistics performed well across our analyses, further investigation may suggest alternative data-driven approaches for choosing the optimal number of PCs.

We choose to use Frobenius and Euclidean distances throughout our analyses; however, other distances could be employed. Several studies have compared choices of distance in various data types including positive semidefinite matrices (Dryden et al., 2009), time series (Wang et al., 2013), and brain connectivity maps (Shehzad et al., 2014). Future investigations could examine how the choices of distance measures influences SiMMR results, particularly when different types of distances (e.g. Euclidean and non-Euclidean) are chosen.

The SiMMR-PC test statistic performs multivariate regression through the principal components of the outcome variable, which is related to previous work in multivariate regression. In particular, SiMMR-PC resembles PC-based test statistics in the multiple phenotype setting with multiple outcome variables and a single covariate (Liu and Lin, 2019); but these statistics do not apply to our settings with multiple covariates. Our investigation is closely related to previous work performing likelihood ratio tests on PCs, which also tested other approaches such as regularization and shrinkage applied to covariance estimates (Ullah and Jones, 2015). Future studies of SiMMR could incorporate other PC-based statistics and alternative high-dimensional test statistics.

## Acknowledgements

This work was supported by the National Institute of Neurological Disorders and Stroke (grant numbers R01 NS085211 and R01 NS060910), the National Multiple Sclerosis Society (RG-1707-28586), the National Institute of Mental Health (R01 MH123550, R01 MH112274, and R01 MH119219), the National Science Foundation Graduate Research Fellowship Program, and a seed grant from the University of Pennsylvania Center for Biomedical Image Computing and Analytics (CBICA). The content is solely the responsibility of the authors and does not necessarily represent the official views of the funding agencies.

## References

Abdi, H., O’Toole, A., Valentin, D., and Edelman, B. (2005). DISTATIS: The Analysis of Multiple Distance Matrices. In 2005 IEEE Computer Society Conference on Computer Vision and Pattern Recognition (CVPR’05) - Workshops, pages 42–42.

Alam, M. A., Qiu, C., Shen, H., Wang, Y.-P., and Deng, H.-W. (2021). A generalized kernel machine approach to identify higher-order composite effects in multi-view datasets, with application to adolescent brain development and osteoporosis. Journal of Biomedical Informatics, 120:103854.

Anderson, M. J. (2001). A new method for non-parametric multivariate analysis of variance. Austral Ecology, 26(1):32–46.

Baum, G. L., Cui, Z., Roalf, D. R., Ciric, R., Betzel, R. F., Larsen, B., Cieslak, M., Cook, P. A., Xia, C. H., Moore, T. M., Ruparel, K., Oathes, D. J., Alexander-Bloch, A. F., Shinohara, R. T., Raznahan, A., Gur, R. E., Gur, R. C., Bassett, D. S., and Satterthwaite, T. D. (2020). Development of structure–function coupling in human brain networks during youth. Proceedings of the National Academy of Sciences, 117(1):771–778.

Cailliez, F. (1983). The analytical solution of the additive constant problem. Psychometrika, 48(2):305–308.

Dale, A. M., Fischl, B., and Sereno, M. I. (1999). Cortical surface-based analysis. I. Segmentation and surface reconstruction. NeuroImage, 9(2):179–194.

Dempster, A. P. (1958). A High Dimensional Two Sample Significance Test. The Annals of Mathematical Statistics, 29(4):995–1010.

Dryden, I. L., Koloydenko, A., and Zhou, D. (2009). Non-Euclidean Statistics for Covariance Matrices, with Applications to Diffusion Tensor Imaging. The Annals of Applied Statistics, 3(3):1102–1123.

Faraway, J. (2014). Regression with Distance Matrices. Journal of Applied Statistics, 41(11):2342–2357.

Fischl, B., Sereno, M. I., and Dale, A. M. (1999). Cortical surface-based analysis. II: Inflation, flattening, and a surface-based coordinate system. NeuroImage, 9(2):195–207.

Gao, J., Li, P., Chen, Z., and Zhang, J. (2020). A Survey on Deep Learning for Multimodal Data Fusion. Neural Computation, 32(5):829–864.

Gretton, A., Borgwardt, K., Rasch, M., Schölkopf, B., and Smola, A. (2007). A Kernel Method for the Two-Sample-Problem. In Advances in Neural Information Processing Systems, volume 19. MIT Press.

Johns, J. T., Di, J., Merikangas, K., Cui, L., Swendsen, J., and Zipunnikov, V. (2019). Fragmentation as a novel measure of stability in normalized trajectories of mood and attention measured by ecological momentary assessment. Psychological Assessment, 31(3):329–339.

Kwee, L. C., Liu, D., Lin, X., Ghosh, D., and Epstein, M. P. (2008). A powerful and flexible multilocus association test for quantitative traits. American Journal of Human Genetics, 82(2):386–397.

Lahat, D., Adali, T., and Jutten, C. (2015). Multimodal Data Fusion: An Overview of Methods, Challenges, and Prospects. Proceedings of the IEEE, 103(9):1449–1477.

Lamers, F., Swendsen, J., Cui, L., Husky, M., Johns, J., Zipunnikov, V., and Merikangas, K. R. (2018). Mood reactivity and affective dynamics in mood and anxiety disorders. Journal of Abnormal Psychology, 127(7):659–669.

Langsrud, Ø. (2004). The geometrical interpretation of statistical tests in multivariate linear regression. Statistical Papers, 45(1):111–122.

Li, J., Zhang, W., Zhang, S., and Li, Q. (2019). A theoretic study of a distance-based regression model. Science China Mathematics, 62(5):979–998.

Liu, Z., Barnett, I., and Lin, X. (2020). A comparison of principal component methods between multiple phenotype regression and multiple SNP regression in genetic association studies. The Annals of Applied Statistics, 14(1):433–451.

Liu, Z. and Lin, X. (2019). A Geometric Perspective on the Power of Principal Component Association Tests in Multiple Phenotype Studies. Journal of the American Statistical Association, 114(527):975–990.

Mardia, K. V., Kent, J. T., and Bibby, J. M. (1979). Multivariate Analysis. Probability and Mathematical Statistics. Academic Press, London ; New York.

McArdle, B. H. and Anderson, M. J. (2001). Fitting Multivariate Models to Community Data: A Comment on Distance-Based Redundancy Analysis. Ecology, 82(1):290–297.

McArtor, D. B., Lubke, G. H., and Bergeman, C. S. (2017). Extending Multivariate Distance Matrix Regression with an Effect Size Measure and the Asymptotic Null Distribution of the Test Statistic. Psychometrika, 82(4):1052–1077.

Merikangas, K. R., Cui, L., Heaton, L., Nakamura, E., Roca, C., Ding, J., Qin, H., Guo, W., Shugart, Y. Y., Yao-Shugart, Y., Zarate, C., and Angst, J. (2014). Independence of familial transmission of mania and depression: Results of the NIMH family study of affective spectrum disorders. Molecular Psychiatry, 19(2):214–219.

Merikangas, K. R., Swendsen, J., Hickie, I. B., Cui, L., Shou, H., Merikangas, A. K., Zhang, J., Lamers, F., Crainiceanu, C., Volkow, N. D., and Zipunnikov, V. (2019). Real-time Mobile Monitoring of the Dynamic Associations Among Motor Activity, Energy, Mood, and Sleep in Adults With Bipolar Disorder. JAMA psychiatry, 76(2):190–198.

Pan, W. (2011). Relationship between genomic distance-based regression and kernel machine regression for multi-marker association testing. Genetic Epidemiology, 35(4):211–216.

Reiss, P. T., Stevens, M. H. H., Shehzad, Z., Petkova, E., and Milham, M. P. (2010). On Distance-Based Permutation Tests for Between-Group Comparisons. Biometrics, 66(2):636–643.

Satterthwaite, T. D., Elliott, M. A., Ruparel, K., Loughead, J., Prabhakaran, K., Calkins, M. E., Hopson, R., Jackson, C., Keefe, J., Riley, M., Mensh, F. D., Sleiman, P., Verma, R., Davatzikos, C., Hakonarson, H., Gur, R. C., and Gur, R. E. (2014). Neuroimaging of the Philadelphia Neurodevelopmental Cohort. NeuroImage, 86:544–553.

Schork, N. J. and Zapala, M. A. (2012). Statistical Properties of Multivariate Distance Matrix Regression for High-Dimensional Data Analysis. Frontiers in Genetics, 3.

Sejdinovic, D., Sriperumbudur, B., Gretton, A., and Fukumizu, K. (2013). Equivalence of Distance-Based and RKHS-Based Statistics in Hypothesis Testing. The Annals of Statistics, 41(5):2263–2291.

Shehzad, Z., Kelly, C., Reiss, P. T., Cameron Craddock, R., Emerson, J. W., McMahon, K., Copland, D. A., Xavier Castellanos, F., and Milham, M. P. (2014). A multivariate distance-based analytic framework for connectome-wide association studies. NeuroImage, 93:74–94.

Shen, C. and Vogelstein, J. T. (2020). The exact equivalence of distance and kernel methods in hypothesis testing. AStA Advances in Statistical Analysis.

Shi, Y., Zhang, W., Liu, A., and Li, Q. (2021). Distance-based regression analysis for measuring associations. 2105.10145 [math, stat].

Shinohara, R. T., Shou, H., Carone, M., Schultz, R., Tunc, B., Parker, D., Martin, M. L., and Verma, R. (2020). Distance-based analysis of variance for brain connectivity. Biometrics, 76(1):257–269.

Shou, H., Cui, L., Hickie, I., Lameira, D., Lamers, F., Zhang, J., Crainiceanu, C., Zipunnikov, V., and Merikangas, K. R. (2017). Dysregulation of objectively assessed 24-hour motor activity patterns as a potential marker for bipolar I disorder: Results of a community-based family study. Translational Psychiatry, 7(8):e1211.

Sudlow, C., Gallacher, J., Allen, N., Beral, V., Burton, P., Danesh, J., Downey, P., Elliott, P., Green, J., Landray, M., Liu, B., Matthews, P., Ong, G., Pell, J., Silman, A., Young, A., Sprosen, T., Peakman, T., and Collins, R. (2015). UK biobank: An open access resource for identifying the causes of a wide range of complex diseases of middle and old age. PLoS medicine, 12(3):e1001779.

Székely, G. J. and Rizzo, M. L. (2014). Partial distance correlation with methods for dissimilarities. The Annals of Statistics, 42(6):2382–2412.

Székely, G. J., Rizzo, M. L., and Bakirov, N. K. (2007). Measuring and testing dependence by correlation of distances. The Annals of Statistics, 35(6):2769–2794.

Ullah, I. and Jones, B. (2015). Regularised Manova for High-Dimensional Data. Australian & New Zealand Journal of Statistics, 57(3):377–389.

Vandekar, S. N., Shinohara, R. T., Raznahan, A., Hopson, R. D., Roalf, D. R., Ruparel, K., Gur, R. C., Gur, R. E., and Satterthwaite, T. D. (2016). Subject-level Measurement of Local Cortical Coupling. NeuroImage, 133:88–97.

Vandekar, S. N., Shinohara, R. T., Raznahan, A., Roalf, D. R., Ross, M., DeLeo, N., Ruparel, K., Verma, R., Wolf, D. H., Gur, R. C., Gur, R. E., and Satterthwaite, T. D. (2015). Topologically Dissociable Patterns of Development of the Human Cerebral Cortex. Journal of Neuroscience, 35(2):599–609.

Wang, X., Mueen, A., Ding, H., Trajcevski, G., Scheuermann, P., and Keogh, E. (2013). Experimental comparison of representation methods and distance measures for time series data. Data Mining and Knowledge Discovery, 26(2):275–309.

Weinstein, S. M., Vandekar, S. N., Adebimpe, A., Tapera, T. M., Robert-Fitzgerald, T., Gur, R. C., Gur, R. E., Raznahan, A., Satterthwaite, T. D., Alexander-Bloch, A. F., and Shinohara, R. T. (2021). A simple permutation-based test of intermodal correspondence. Human Brain Mapping, 42(16):5175–5187.

Zhao, N., Chen, J., Carroll, I. M., Ringel-Kulka, T., Epstein, M. P., Zhou, H., Zhou, J. J., Ringel, Y., Li, H., and Wu, M. C. (2015). Testing in Microbiome-Profiling Studies with MiRKAT, the Microbiome Regression-Based Kernel Association Test. American Journal of Human Genetics, 96(5):797–807.

