## Supplementary materials for "Similarity-Based Multimodal Regression"

### Similarity-Based Multimodal Regression Supplementary Materials

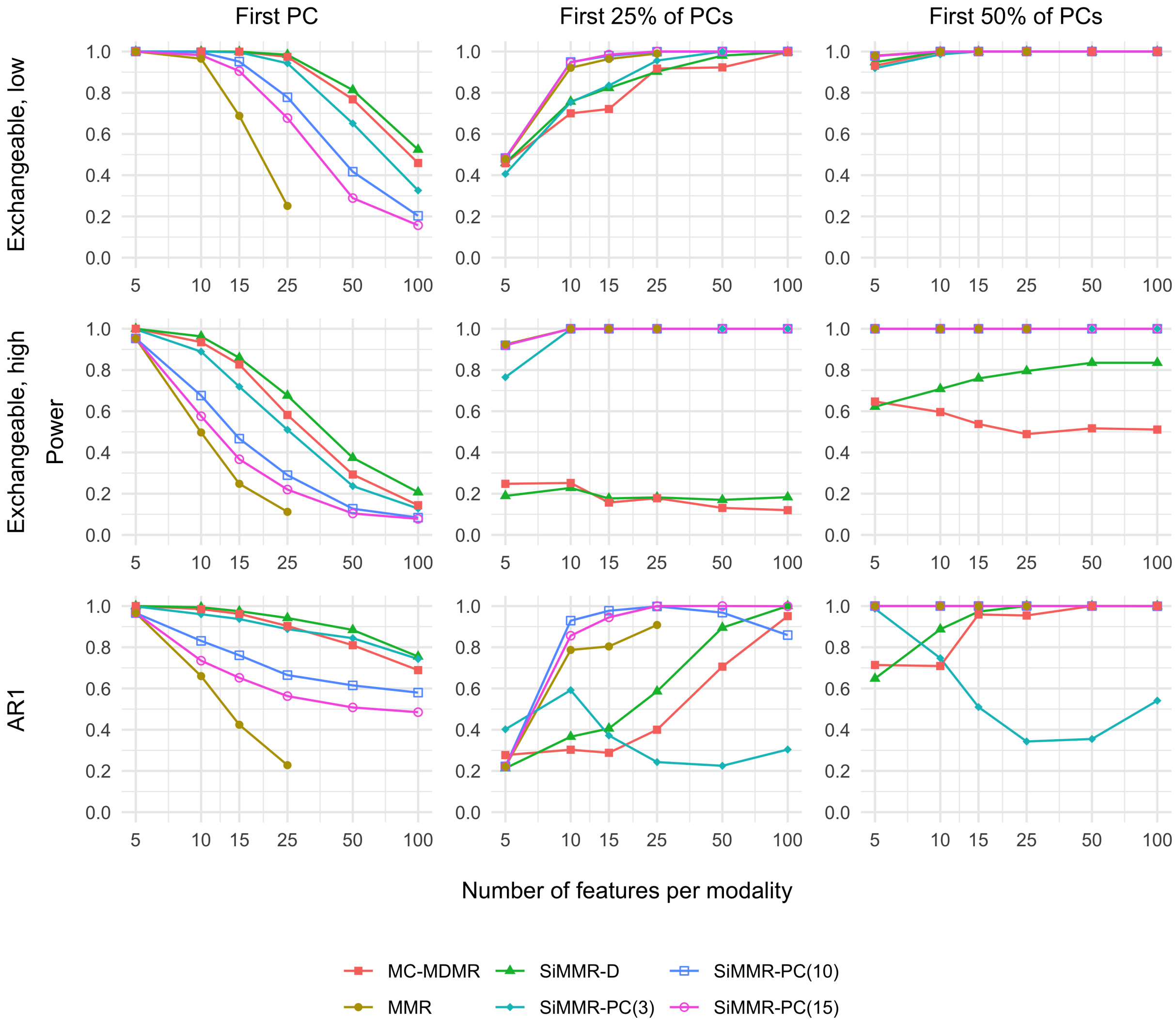


Supplementary Figure 1: **Results in simulations with exchangeable and AR(1) correlation structures for a sample size of 100.** Each trace represents a different test statistic. Different simulation settings are distinguished by correlation structure across rows and by rank of the binary covariate effect across columns. Abbreviations: MDMR, multivariate distance matrix regression; MC-MDMR, multiple MDMR statistics after Bonferroni correction; MMR, multivariate multiple regression using Pillai's trace.

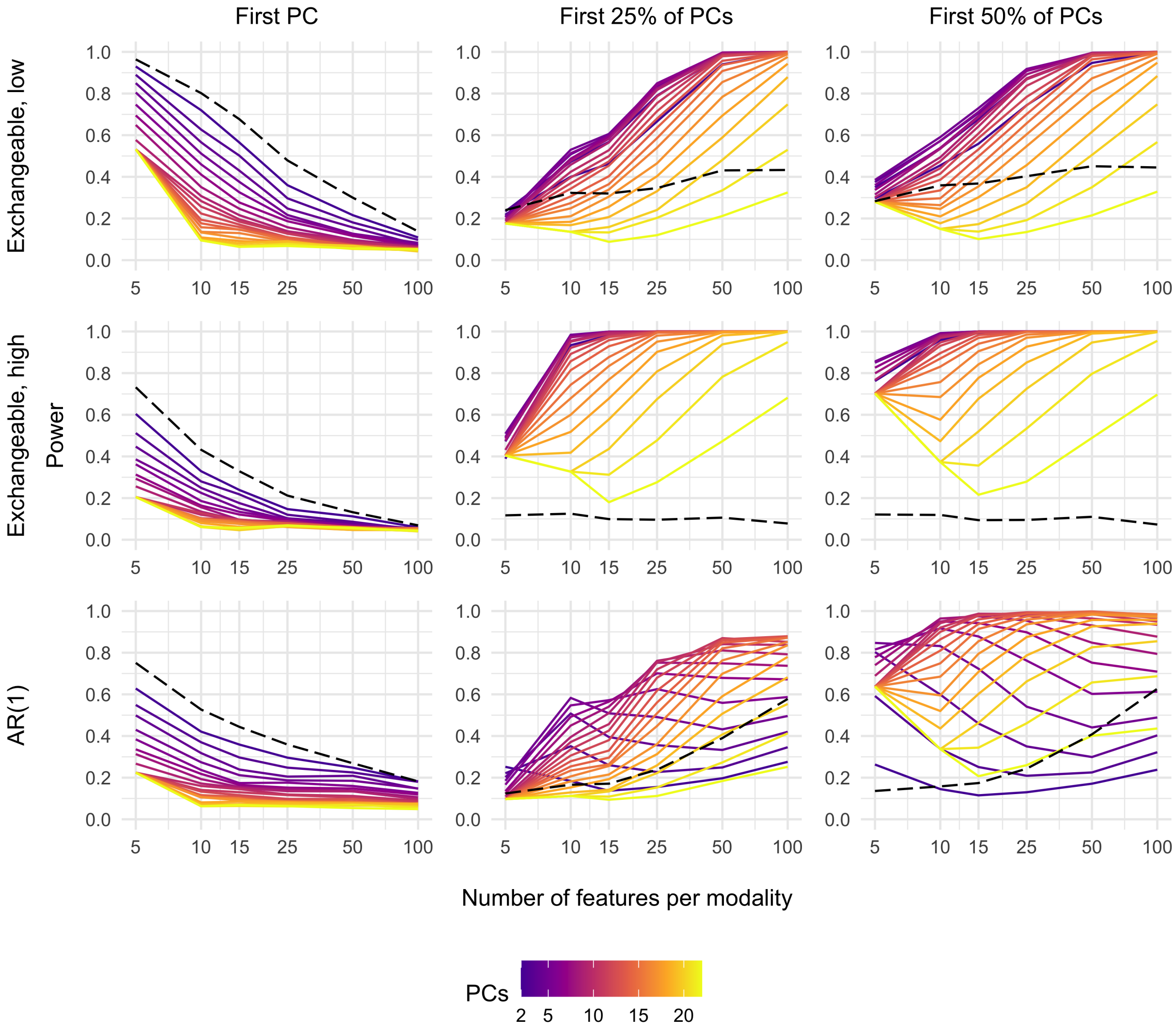


Supplementary Figure 2: **SiMMR-PC power results in simulations with exchangeable and AR1 correlation structures with a sample size of 25.** Results are shown for SiMMR-PC with PCs ranging from 2 to 25. The dashed line represents the power of SiMMR-D for comparison.

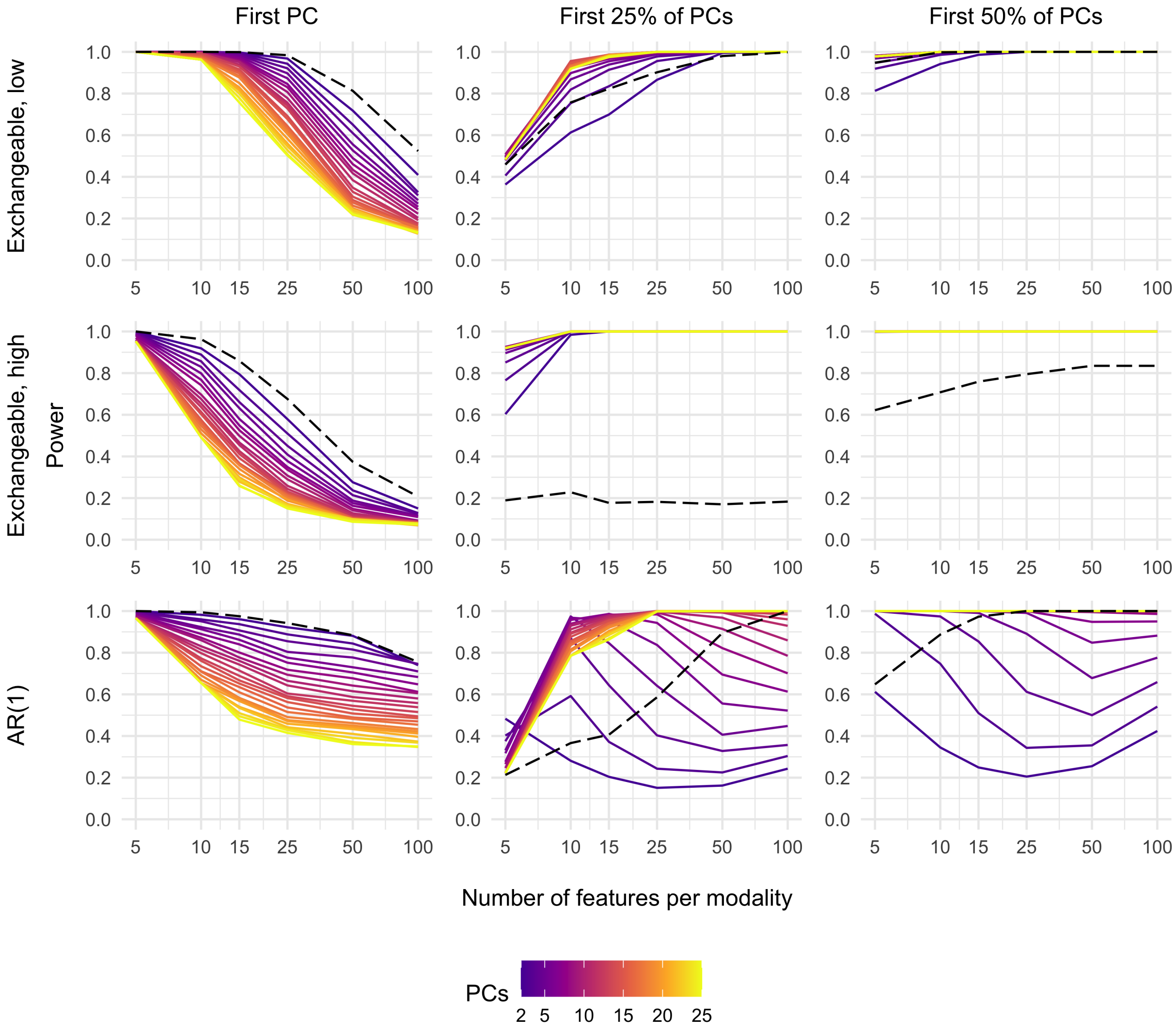


Supplementary Figure 3: **SiMMR-PC power results in simulations with exchangeable and AR1 correlation structures with a sample size of 100.** Results are shown for SiMMR-PC with PCs ranging from 2 to 25. The dashed line represents the power of SiMMR-D for comparison.

| PC | $N$ | $M$ | $Q$ | MMR | MC-MDMR | SiMMR-D | SiMMR-PC(3) | SiMMR-PC(10) | SiMMR-PC(15) |
|---|---|---|---|---|---|---|---|---|---|
| None | 25 | 2 | 5 | 0.054 | 0.042 | 0.048 | 0.051 | 0.052 | 0.052 |
| | | | 25 | | 0.048 | 0.044 | 0.051 | 0.049 | 0.045 |
| | | | 100 | | 0.036 | 0.037 | 0.042 | 0.038 | 0.045 |
| | | 10 | 5 | | 0.032 | 0.048 | 0.033 | 0.035 | 0.037 |
| | | | 25 | | 0.045 | 0.044 | 0.037 | 0.053 | 0.056 |
| | | | 100 | | 0.05 | 0.041 | 0.04 | 0.041 | 0.054 |
| | 100 | 2 | 5 | 0.051 | 0.052 | 0.055 | 0.05 | 0.048 | 0.048 |
| | | | 25 | 0.053 | 0.045 | 0.042 | 0.047 | 0.048 | 0.046 |
| | | | 100 | | 0.044 | 0.046 | 0.038 | 0.048 | 0.052 |
| | | 10 | 5 | 0.04 | 0.029 | 0.05 | 0.06 | 0.039 | 0.048 |
| | | | 25 | | 0.051 | 0.042 | 0.042 | 0.044 | 0.042 |
| | | | 100 | | 0.051 | 0.055 | 0.045 | 0.048 | 0.05 |
| 25% | 25 | 2 | 5 | 0.094 | 0.146 | 0.124 | **0.221** | 0.097 | 0.097 |
| | | | 25 | | 0.184 | 0.238 | 0.227 | **0.754** | 0.509 |
| | | | 100 | | 0.396 | 0.578 | 0.346 | 0.837 | **0.872** |
| | | 10 | 5 | | 0.155 | 0.219 | 0.202 | **0.748** | 0.511 |
| | | | 25 | | 0.275 | 0.733 | 0.428 | 0.824 | **0.882** |
| | | | 100 | | 0.665 | **1** | 0.935 | 0.995 | 0.989 |
| | 100 | 2 | 5 | 0.221 | 0.277 | 0.213 | **0.402** | 0.222 | 0.222 |
| | | | 25 | 0.908 | 0.4 | 0.586 | 0.243 | 0.998 | **1** |
| | | | 100 | | 0.951 | **1** | 0.304 | 0.859 | 0.999 |
| | | 10 | 5 | 0.9 | 0.409 | 0.568 | 0.222 | **1** | **1** |
| | | | 25 | | 0.726 | **1** | 0.344 | 0.834 | 0.99 |
| | | | 100 | | 0.999 | **1** | 0.923 | **1** | **1** |
| 50% | 25 | 2 | 5 | **0.64** | 0.123 | 0.136 | 0.592 | 0.637 | 0.637 |
| | | | 25 | | 0.173 | 0.243 | 0.209 | **0.986** | 0.974 |
| | | | 100 | | 0.4 | 0.627 | 0.322 | 0.934 | **0.984** |
| | | 10 | 5 | | 0.149 | 0.227 | 0.179 | **0.986** | **0.986** |
| | | | 25 | | 0.205 | 0.804 | 0.434 | 0.903 | **0.979** |
| | | | 100 | | 0.528 | **1** | 0.948 | **1** | 0.995 |
| | 100 | 2 | 5 | **1** | 0.714 | 0.648 | 0.987 | **1** | **1** |
| | | | 25 | **1** | 0.954 | **1** | 0.343 | **1** | **1** |
| | | | 100 | | **1** | **1** | 0.541 | 0.999 | **1** |
| | | 10 | 5 | **1** | 0.926 | **1** | 0.317 | **1** | **1** |
| | | | 25 | | **1** | **1** | 0.67 | **1** | **1** |
| | | | 100 | | **1** | **1** | **1** | **1** | **1** |

Supplementary Table 1: **Simulation results for the AR(1) correlation setting.** Rejection rates are shown across varying number of subjects (N), number of modalities (M), number of features per modality (Q), and number of principal components included in the binary covariate effect (PC). The highest power in each row is bolded.
